# Investigating genetic correlations and causal effects between caffeine consumption and sleep behaviours

**DOI:** 10.1101/199828

**Authors:** Jorien L. Treur, Mark Gibson, Amy E Taylor, Peter J Rogers, Marcus R Munafò

**Author notes:** Corresponding author: Jorien L Treur, School of Experimental Psychology, University of Bristol, 12a Priory Road, Bristol BS8 1TU, United Kingdom.

## Abstract

**Study Objectives:** Higher caffeine consumption has been linked to poorer sleep and insomnia complaints. We investigated whether these observational associations are the result of genetic risk factors influencing both caffeine consumption and poorer sleep, and/or whether they reflect (possibly bidirectional) causal effects. **Methods:** Summary-level data were available from genome-wide association studies (GWAS) on caffeine consumption (n=91,462), sleep duration, and chronotype (i.e., being a ‘morning’ versus an ‘evening’ person) (both n=128,266), and insomnia complaints (n=113,006). Linkage disequilibrium (LD) score regression was used to calculate genetic correlations, reflecting the extent to which genetic variants influencing caffeine consumption and sleep behaviours overlap. Causal effects were tested with bidirectional, two-sample Mendelian randomization (MR), an instrumental variable approach that utilizes genetic variants robustly associated with an exposure variable as an instrument to test causal effects. Estimates from individual genetic variants were combined using inverse-variance weighted meta-analysis, weighted median regression and MR Egger regression methods. **Results:** There was no clear evidence for genetic correlation between caffeine consumption and sleep duration (*r*g=0.000, *p*=0.998), chronotype (*r*g=0.086, *p*=0.192) or insomnia (*r*g=-0.034, *p*=0.700). Two-sample Mendelian randomization analyses did not support causal effects from caffeine consumption to sleep behaviours, or the other way around. **Conclusions:** We found no evidence in support of genetic correlation or causal effects between caffeine consumption and sleep. While caffeine may have acute effects on sleep when taken shortly before habitual bedtime, our findings suggest that a more sustained pattern of high caffeine consumption is likely associated with poorer sleep through shared environmental factors.

## Introduction

Caffeine is the most commonly used psychoactive substance, with coffee being the second most popular beverage worldwide (after water) (1). There are also cultural differences in the popularity of caffeinated beverages, with tea being more popular than coffee in some countries such as the United Kingdom (2). Acutely, caffeine is known to affect alertness and concentration through its antagonistic effects on adenosine receptors (3, 4), although because of tolerance the net benefit of frequent caffeine consumption appears to be negligible (5). Consumption of caffeinated beverages has also been linked to poor sleep – a recent review of the literature showed that higher caffeine consumption is associated with prolonged sleep latency (the time it takes to fall asleep), reduced sleep time, reduced sleep efficiency (percentage of time asleep of the total time in bed), and poorer sleep quality (6). Moreover, caffeine consumption correlates positively with insomnia complaints (7, 8) and negatively with chronotype (being a ‘morning’ versus an ‘evening’ person) (9, 10). Given the higher mortality rates and poorer health outcomes associated with sleep problems (11, 12), it is important to understand how caffeine consumption relates to different sleep behaviours.

The co-occurrence of high caffeine consumption and poor sleep may be the result of different (not mutually exclusive) mechanisms. Factors that increase a person’s caffeine consumption may also increase their risk of sleeping problems. Such overlapping risk factors could be environmental in nature, or genetic. However, there may (also) be causal effects between caffeine consumption and sleep behaviours. Given the well-known stimulating effects of caffeine, which is often the reason people consume caffeinated beverages, it is plausible that sustained high caffeine consumption causes problems with sleeping. In extreme cases, it may even cause or exacerbate symptoms of insomnia. Controlled laboratory studies suggest that caffeine impacts human sleep quality. However, these studies typically administer caffeine shortly before habitual bedtime (i.e., ≤60 minutes before), which may not reflect real life consumption patterns. In addition, most studies to date have been conducted in male participants only (6) and causal effects in the other direction have not been tested – individuals who tend to sleep less and/or have insomnia may consume more caffeine to alleviate the effects of sleep deprivation during the day. Novel methods are needed to fully disentangle possible causal mechanisms.

To determine whether observational associations between caffeine consumption and sleep variables are due to overlapping genetic risk factors and/or causal effects (in either direction), we applied two methods. First, we calculated genetic correlations between caffeine consumption and sleep duration, insomnia complaints and chronotype based on summary level data of recent large-scale genome-wide association studies (GWAS) (13-15). While the outcome of the GWAS on caffeine consumption was cups of coffee consumed per day, genetic risk scores composed of the top genetic hits have been shown to associate more generally with other types of caffeinated beverages (e.g., tea) as well (16). Genetic correlations reflect the extent to which genetic variants that influence caffeine consumption also influence sleep behaviours. Evidence of genetic correlation indicates shared genetic aetiologies but may also (partly) reflect causal effects. To further investigate possible causal effects and their direction, we also applied two sample Mendelian randomization (MR) analysis. This instrumental variable approach utilizes a selection of genetic variants that are robustly associated with an exposure variable as an instrument to test causal effects on an outcome variable. Potential biological pleiotropy (i.e., effects of genotype on the outcome of interest not acting through the exposure) can be tested with sensitivity analyses. By combining two novel research methods we aim to disentangle mechanisms underlying observational associations between caffeine consumption and sleep behaviours.

## Methods

### Study population

For caffeine consumption, we used summary statistics from the Coffee and Caffeine Genetics Consortium GWAS (n=91,462) (13). For sleep behaviours, GWAS summary statistics were available for sleep duration in hours of sleep and chronotype (a continuous score of being a ‘morning’ versus an ‘evening’ person) (n=128,266) (14), and for insomnia (usually having trouble falling asleep at night or waking up in the middle of the night (‘cases’) versus never/rarely or sometimes having these problems (‘controls’)) (n=113,006) (15). The GWAS on sleep behaviours were performed in UK Biobank and there was no sample overlap with the GWAS on caffeine consumption.

### LD score regression

The main premise of LD score regression is that genetic variants that are in high linkage disequilibrium (LD) with other genetic variants across the genome, are more likely to tag a causal genetic variant – one that exerts a true, causal effect on the phenotype in question – than genetic variants that are in low LD with other genetic variants. Based on this expected relationship between LD and the strength of association, for two phenotypes, a genetic correlation can be calculated. Genetic correlation reflects to what degree the genetic liability for one phenotype correlates with the genetic liability for a second phenotype. LD score regression methods have been described in more detail previously (17). We calculated genetic correlations using the summary data described above. Pre-calculated and publicly-available LD scores – the degree of LD a single nucleotide polymorphism (SNP) has with all neighbouring SNPs – based on individuals of European ancestry were retrieved from https://github.com/bulik/ldsc.

### Mendelian randomization

Mendelian randomisation (MR) uses genetic variants that are robustly associated with an exposure variable as an instrument to test causal effects on an outcome variable (18, 19). With conventional epidemiological methods, it is difficult to determine the causality because an observational association can also be the result of confounding factors that predict both variables (e.g., socio-economic position) or reverse causality (an outcome variable affecting the exposure variable). MR is in principle better protected against confounding than conventional epidemiological methods because genetic variants are randomly transmitted in the population. Additionally, reverse causality cannot affect MR results because an outcome variable cannot change a person’s genotype. There are three important assumptions to MR: 1) the genetic instrument should be robustly associated with the exposure variable, 2) the genetic instrument should be independent of confounders, and 3) there should be no biological (or horizontal) pleiotropy, meaning that the genetic instrument should not affect the outcome variable through an independent pathway, other than through its effect on the exposure variable.

Here, we applied *two-sample* MR, in which a genetic instrument is first identified in a GWAS of the exposure variable and then the same instrument is identified in a second, separate GWAS of the outcome variable (20). When the genetic instrument was composed of a single genetic variant the Wald ratio method was applied (gene-outcome association / gene-exposure association; (21)) when the instrument comprised multiple genetic variants, the ratios were combined in an inverse-variance weighted (IVW) meta-analysis (summing ratio estimates of all variants in a weighted average formula; (22)). To test the third MR assumption (no horizontal pleiotropy) we additionally used two sensitivity analyses. First, we used the weighted median approach, which is a method that can provide a consistent estimate of a causal effect even in a situation where up to 50% of the weight comes from invalid instruments (23). Second, we used MR-Egger regression, which applies Egger’s test, normally used to assess small study bias in meta-analyses, to polygenic MR instruments (22).Under MR-Egger it is assumed that there is no correlation between the strength of an instrument (SNP-exposure association) and the effect that the instrument has on the outcome. This is referred to as the InSIDE assumption (Instrument Strength Independent of Direct Effect) and it is a much weaker assumption than the assumption of no horizontal pleiotropy (22).

Genetic instruments were first identified for caffeine consumption after which causal effects on sleep behaviours (sleep duration, chronotype and insomnia) were tested. Next, genetic instruments for the different sleep behaviours were identified and causal effects on caffeine consumption were tested. For each phenotype, we constructed two genetic instruments; one consisting of SNPs that were associated with the exposure variable under the genome-wide significant *p*-value threshold of *p*<5×10^-8^ and one consisting of SNPs associated with the exposure variable under a more lenient p-value threshold of *p*<1×10^-5^. All analyses were performed in the database and analytical platform *MR-Base* (24). For instruments of threshold *p*<5×10^-8^, all independent genome-wide significant hits were selected manually from the published GWAS papers (based on the discovery samples) and then introduced to MR-Base while instruments of threshold *p*<1×10^-5^ were constructed in MR-Base (including the pruning of genetic variants (*r^2^*<0.001) and retrieving of proxies (*r*^*2*^≥0.8)). Details on the SNPs included in all genetic instruments are provided in Supplementary Table 1.

**Table 1.**
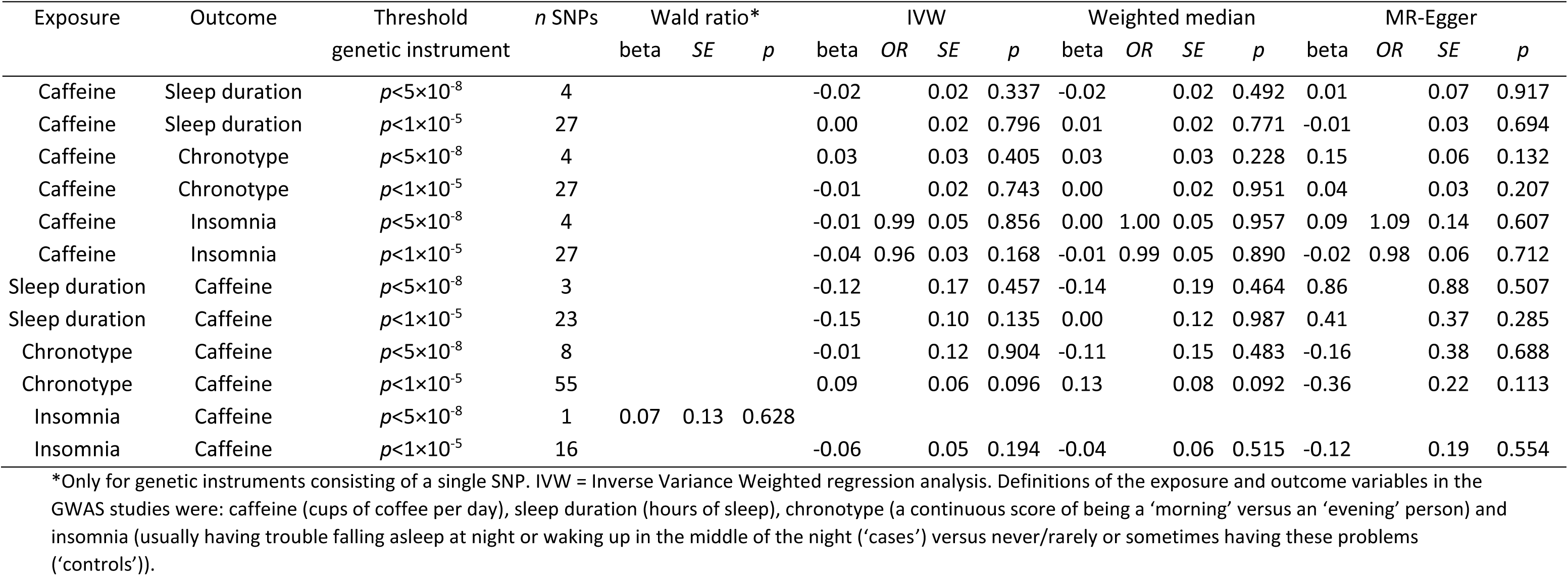
Bidirectional, two sample Mendelian randomization analyses between caffeine consumption and sleep behaviours.

## Results

With LD score regression, we found no clear evidence for genetic correlation between sleep duration and caffeine consumption (*r*g=0.000, SE=0.079, *p*=0.998), chronotype and caffeine consumption (*r*g=0.086, SE=0.066, *p*=0.192), or insomnia and caffeine consumption (*r*g=-0.034, SE=0.087, *p*=0.700).

Two sample MR, using all three analytical approaches, did not provide clear evidence for causal effects of caffeine consumption on sleep duration (IVW beta estimates were -0.02, *p*=0.337 and -0.00, *p*=0.796 for genetic instruments with threshold *p*<5×10^-8^ and *p*<1×10^-5^, respectively), chronotype (beta=0.03, *p*=0.405 and beta=-0.01, *p*=0.743, respectively) or insomnia (beta=-0.01, *p*=0.856 and beta=-0.04, *p*=0.168), respectively). There was also no clear evidence for causal effects from sleep duration to caffeine consumption (beta=-0.12, *p*=0.457 and beta=-0.15. *p*=0.135, respectively), chronotype to caffeine consumption (beta=-0.01, *p*=0.904 and beta=0.09, *p*=0.096) or insomnia to caffeine consumption (beta=0.07, *p*=0.628 and beta=-0.06. *p*=0.194, respectively). More details of these results are shown in **Table 1**. Cochran’s heterogeneity statistic (Q), which assesses heterogeneity between the different SNPs included in a genetic instrument, indicated heterogeneity for IVW analyses from caffeine consumption to chronotype (see Supplementary Table 2). The intercepts from MR-Egger regression analyses, which estimate the degree of biological pleiotropy, did not provide strong evidence for pleiotropy overall, although there was some weak evidence for pleiotropy from chronotype to caffeine consumption (see Supplementary Table 3).

## Discussion

We did not find clear evidence in support of a genetic correlation between caffeine consumption on the one hand and sleep duration, insomnia or chronotype on the other hand. In addition, our findings from Mendelian randomisation analyses did not support causal relationships from caffeine consumption to sleep behaviours, or the other way around. Together, these results suggest that previously reported, observational associations between high caffeine consumption and poor sleep are likely due to environmental factors that influence both.

Our findings corroborate previous reports showing that none of the genetic variants associated with caffeine consumption were associated with caffeine-induced insomnia (13, 25). In contrast, controlled laboratory studies have suggested that caffeine has a causal, negative impact on sleep (6). It should be noted though, that in most of these studies, participants were administered caffeine acutely, immediately before their usual bedtime. In the current study, we measured genetic liability for caffeine consumption, a measure that reflects a more sustained life-time average intake of caffeine, and not only intake just before going to sleep. It may be the case that caffeine impacts sleep when it is consumed in the evening, while there is little or no effect when it is consumed during the day. It is likely that most caffeine is consumed earlier during the day, given that a common reason for consuming caffeinated beverages is their stimulant effects (26, 27). One small study (n=12) looked at the effects of a high dose of caffeine (400 mg, similar to the amount of caffeine in at least 4 cups of coffee) on sleep when administered 0, 3 or 6 hours before bedtime and did find disruptive effects on sleep at all time points (28). Another possibility for the lack of evidence for causal effects in the present study is that, over time, tolerance to the effects of caffeine develops (5). In other words, frequent consumption of caffeine does in itself not disrupt sleep, while caffeine withdrawal actually increases sleepiness – this is what has been observed for daytime sleepiness (5). While the genetic instrument that we used for caffeine has been robustly associated with higher caffeine consumption (13), these same genetic variants are also associated with lower circulating caffeine levels (32). The most likely explanation for this is that even though these individuals consume more caffeine, they metabolise caffeine more rapidly and therefore show lower blood concentrations of caffeine and its metabolites. This could also mean that, despite influencing higher consumption, faster metabolism of caffeine means that these variants have negligible effects on sleep. Finally, while many of the previous laboratory studies included male participants only, our findings are based on very large samples of males and females (of European ancestry).

In the direction from sleep behaviours to caffeine, we also did not find evidence for causal effects. This is in contrast to research showing that a common reason for changing coffee consumption is experiencing sleep problems (29). It may be that such causal effects did not emerge in our analyses because these are only short-term adjustments in caffeine use that do not hold in the longer term, while our genetic approach reflects a longer-term measure of caffeine consumption.

The lack of evidence for genetic correlation between caffeine consumption and sleep behaviours, and for causal effects, suggests that observational associations may be the result of shared environmental factors. The literature on this topic is scarce, but an example of an environmental factor that could be responsible for both increasing caffeine consumption and inducing or exacerbating sleeping problems is work or school-related demands and stress (30, 31). Daily stress may cause people to have trouble sleeping and may consequently cause them to attempt to self-medicate by consuming more caffeine. More research is needed to identify the environmental factors that increase both caffeine consumption and sleeping problems, to guide the development of more evidence-based interventions to improve sleep.

A major strength to our approach, using summary-level data of very large sample sizes, is that it provides much power to detect small effects which are likely for complex traits such as caffeine consumption and sleeping behaviours. A limitation to our study is that for the Mendelian randomization analyses, we assumed the caffeine consumption SNPs to be associated with caffeine in the GWAS of the sleeping variables, but we were not able to test this. The genetic instrument may be weaker if the GWAS of the outcome variable contains a group of people that do not consume coffee. However, we have previously shown that the genetic risk score of caffeine consumption also predicts coffee consumption in the combined sample of coffee and non-coffee drinkers in UK Biobank (15).

In summary, we did not find clear evidence for causal effects from caffeine consumption to sleep behaviours, or vice versa. Our findings highlight the complexity of interpreting Mendelian randomization results for health behaviours such as caffeine consumption and sleep. While there are well-known acute effects of caffeine on alertness this did not translate to evidence for causal effects of a more sustained intake of caffeine on sleep. Researchers applying Mendelian randomization should be aware that genetic variants used as an instrument, or proxy, for an (exposure) variable, reflect a lifetime-exposure to higher or lower levels of that variable.

## Acknowledgements

AET and MRM are members of the UK Centre for Tobacco and Alcohol Studies, a UKCRC Public Health Research: Centre of Excellence. Funding from British Heart Foundation, Cancer Research UK, Economic and Social Research Council, Medical Research Council, and the National Institute for Health Research, under the auspices of the UK Clinical Research Collaboration, is gratefully acknowledged. JLT is supported by a Rubicon grant from the Netherlands Organization for Scientific Research (NWO; grant number 446-16-009).

